# Small extracellular vesicles promote stiffness-mediated metastasis

**DOI:** 10.1101/2023.07.01.545937

**Authors:** Alexandra Sneider, Ying Liu, Bartholomew Starich, Wenxuan Du, Carolyn Marar, Najwa Faqih, Gabrielle E. Ciotti, Joo Ho Kim, Sejal Krishnan, Salma Ibrahim, Muna Igboko, Alexus Locke, Daniel M. Lewis, Hanna Hong, Michelle Karl, Raghav Vij, Gabriella C. Russo, Praful Nair, Estibaliz Gómez-de-Mariscal, Mehran Habibi, Arrate Muñoz-Barrutia, Luo Gu, T.S. Karin Eisinger-Mathason, Denis Wirtz

## Abstract

Tissue stiffness is a critical prognostic factor in breast cancer and is associated with metastatic progression. Here we show an alternative and complementary hypothesis of tumor progression whereby physiological matrix stiffness affects the quantity and protein cargo of small EVs produced by cancer cells, which in turn drive their metastasis. Primary patient breast tissue produces significantly more EVs from stiff tumor tissue than soft tumor adjacent tissue. EVs released by cancer cells on matrices that model human breast tumors (25 kPa; stiff EVs) feature increased adhesion molecule presentation (ITGα_2_β_1_, ITGα_6_β_4_, ITGα_6_β_1_, CD44) compared to EVs from softer normal tissue (0.5 kPa; soft EVs), which facilitates their binding to extracellular matrix (ECM) protein collagen IV, and a 3-fold increase in homing ability to distant organs in mice. In a zebrafish xenograft model, stiff EVs aid cancer cell dissemination through enhanced chemotaxis. Moreover, normal, resident lung fibroblasts treated with stiff and soft EVs change their gene expression profiles to adopt a cancer associated fibroblast (CAF) phenotype. These findings show that EV quantity, cargo, and function depend heavily on the mechanical properties of the extracellular microenvironment.

**Graphical Abstract:** 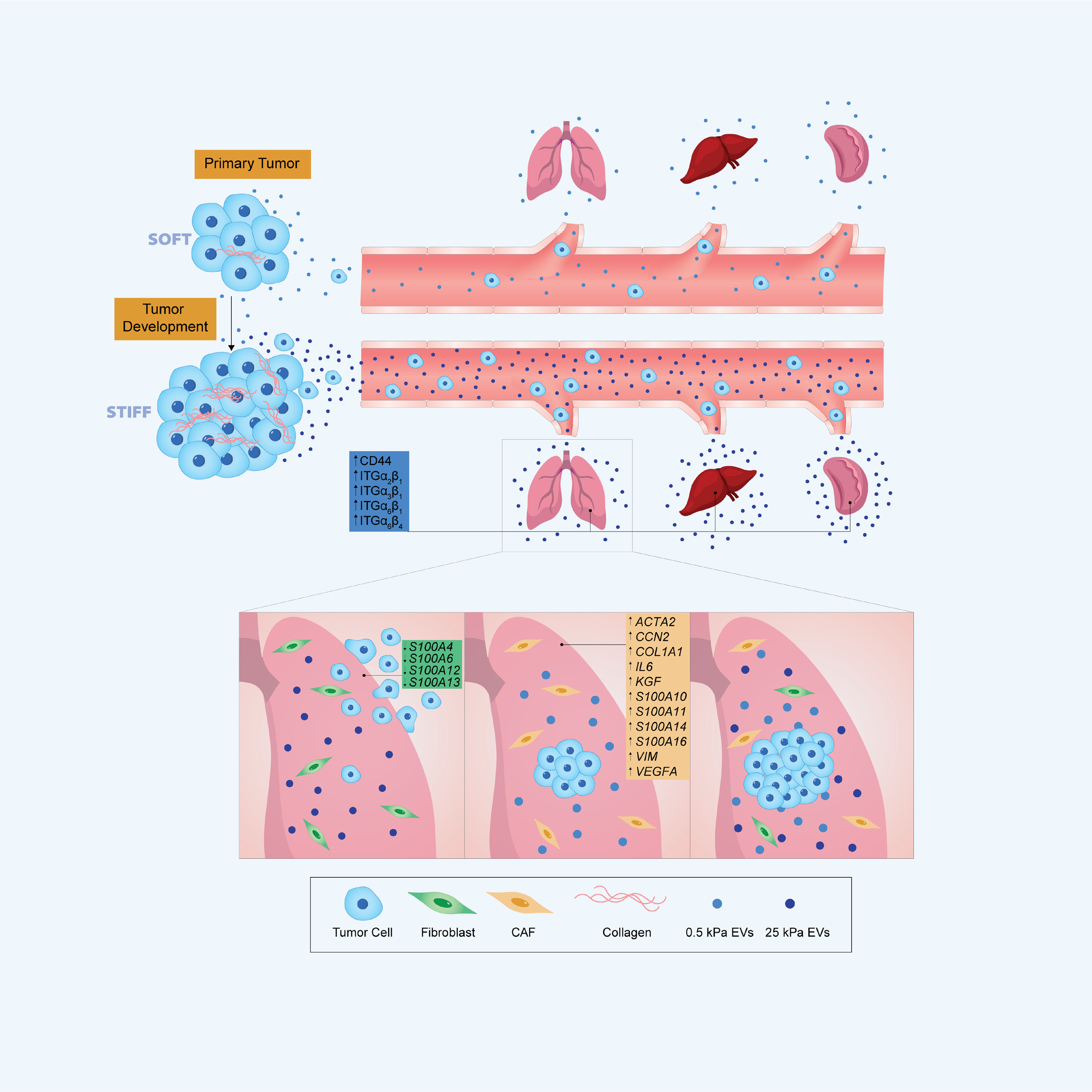

## Introduction

The extracellular matrix (ECM), a network of acellular components predominantly made of collagen, controls tissue structure, modulates cell adhesion, influences secretome dissemination, and conveys mechanical signals^1–4^. Tissues inherently have unique structures and stiffnesses that lend to specific biological processes^5–9^. An increase in ECM stiffness often correlates with poor prognosis in solid tumors^10–15^, explained in part by stiffness-mediated enhanced cancer cell migration and proliferation at the primary tumor^16–19^. As tumors develop, the density and composition of the ECM changes^20^. Due to chronic inflammation, often fibrotic tissue forms at and around the tumor site^21–23^. The increased cross-linking of the ECM can lead to leaky vasculature and promote intravasation^24–27^. Cellular phenotypes also change to promote tumor progression, i.e., fibroblasts develop a cancer associated phenotype that increases the deposition of fibrillar collagen and pro-tumorigenic signaling^28,29^. Despite knowing the mechanical complexities of tumor growth and metastasis, cancer research and the development of therapeutics relies heavily on static model systems, like tissue-culture plastic, that do not incorporate physiologically relevant parameters^30,31^.

Since their discovery, extracellular vesicles (EVs) have primarily been collected and analyzed from tumor cells grown on tissue culture-treated plastic ware. The physiological relevance of EVs obtained in this manner is unknown. EVs are a diverse group of lipid bilayer encapsulated particles secreted by cells that display and encapsulate functional proteins and nucleic acids^32,33^. Small EVs, particularly exosomes that are between 30-150 nm in diameter, have shown great promise as biomarkers and therapeutic agents for the treatment of disease^34–38^. Due to their size, exosomes have the potential to disseminate great distances from their site of secretion^39,40^. Small EVs can transfer their cargo to other cell types and influence homeostasis and disease progression^36,41–47^. Cancer-derived exosomes can increase vascular leakiness, reprogram bone marrow progenitors, increase tumor growth and metastasis^48^; facilitate pre-metastatic niche formation^49–51^; more effectively fuse with target cells^52^; and aid in evading immune detection^53^. Therefore, we sought to characterize the importance of physiologically relevant physical tissue properties (e.g., stiffness) in EV-mediated metastatic dissemination.

Herein we explore how modulating stiffness in the tumor microenvironment can have far-reaching implications on cancer progression through EVs. We observed in primary patient tissue that more EVs are secreted from stiff tissue than softer tissue. We investigated exosomes isolated from breast cancer cells cultured on plastic, 25 kPa (breast tumor stiffness, stiff), and 0.5 kPa (normal tissue stiffness, soft) substrates. EVs from cells on substrates at tumor tissue stiffness have different cargo than those vesicles from soft and plastic substrates. The stiff EV cargo is enriched in integrins (ITGα_2_β_1_, ITGα_6_β_4_, ITGα_6_β_1_), adhesion proteins (CD44), and immune evasion signals over the soft and plastic EVs. These stiff EVs are better able to reach and be retained in distant tissues *in vivo* in mice and adhere to specific ECM proteins like Collagen IV. Additionally, the stiff EVs promote cancer cell motility *in vitro* through a transwell assay, as well as dissemination *in vivo* in zebrafish over soft EVs. EVs isolated from cells cultured on plastic do not consistently match either the physiological stiff or soft conditions in any of the functional assays. Once cancer cells have arrived in distant tissues, the cells experience the mechanically soft environment of normal tissue. While stiff EVs appear to downregulate immune signaling from resident fibroblasts in the lung via a decrease in expression in *S1004*, *S1006*, *S10012*, and *S10013*, potentially to aid cancer cells in evading immune detection, the soft EVs demonstrate the ability to upregulate expression of CAF markers (*ACTA2*, *COL1A1*, *VEGFA*) and inflammatory signals (*S10010*, *S10011*, *S10014*, and *S10016*) in the resident fibroblasts. These results suggest that matrix stiffness influences vesicular secretion and cargo to aid cancer cells at different stages of the metastatic cascade.

## Results

### Physiologically relevant tissue stiffness impacts EV secretion in patients

To determine physiologically relevant stiffnesses, we obtained primary patient breast tumor and adjacent normal tissues for mechanical measurements. Using the compression test, a method that utilizes uniaxial compression, we found a statistically significant difference in the mean Young’s modulus of tumor tissues (19.9 ± 7.1 kPa) and tumor adjacent tissues (2.4 ± 0.5 kPa) (Fig. 1A). Tumor tissue stiffness was further mapped using microindentation, a method that determines the local elastic modulus of evenly spaced points (Fig. S1A and S1B). To investigate the effect of tumor stiffness on EVs, we separated the stiff sections (24.4 ± 4.4 kPa, mean ± SEM) and the soft sections (5.7 ± 0.4 kPa) of the tumor tissues based on the microindentation results. We noted significant intra-tumoral and inter-tumoral heterogeneity, ranging from 2.9 to 81.7 kPa (Fig. S1A). Based on these findings we elected to use a 25 kPa matrix to represent stiffer human tumor tissue in our subsequent assays. Given our interest in investigating the impact of EVs at distant sites, such as the lung which has a stiffness ranging from 0.5-5 kPa^48,54–59^, we chose a matrix stiffness of 0.5 kPa to represent softer tissues. We compared EVs collected from cells grown on matrices at these physiological stiffnesses to EVs derived from cells grown on plastic culture dishes with non-physiological stiffness between 2 and 4 GPa^7^.

**Figure 1:**
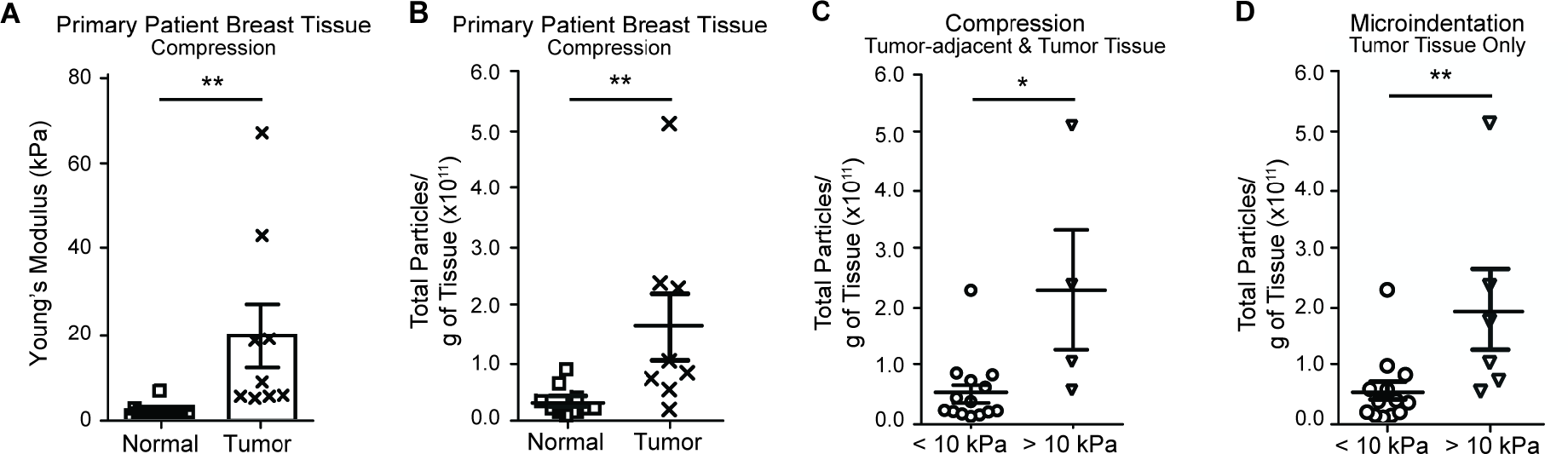
Matrix stiffness impacts the quantity of EVs produced by patient tissue. **A**, Mean elastic moduli (kPa, mean ± SEM) of patient tumor and adjacent normal tissues by compression measurements. Ten adjacent normal tissue samples and nine tumor tissue samples. Non-parametric t-test. Vesicles released per gram of tissue in patient breast tissue samples: **B,** Pathologist determined tumor-adjacent or tumor tissue. Ten normal tissue samples and eight tumor tissue samples. **C**, Tumor-adjacent, and tumor tissues pooled together. Samples separated by mean compression measurements (kPa, mean ± SEM). Fourteen samples < 10 kPa and four samples > 10 kPa. **D**, Tumor samples separated by mean microindentation measurements (kPa, mean ± SEM). Thirteen samples < 10 kPa and six samples > 10 kPa. Non-parametric t-test.

Following microindentation analyses of resected human breast cancer samples, we sectioned the tissues by stiffness and isolated EVs from stiff and soft regions. To preserve the integrity and micromechanics of these tissues, the tissues were not dissociated; therefore, isolated vesicles were released from both cancer and tumor-associated cells. Using compression analysis, tumor samples released significantly more vesicles per gram of tissue than tumor adjacent tissue (Fig. 1B, and Fig. S1C). Significantly more vesicles were released per gram of tissue with a mean tissue stiffness > 10 kPa than from tissues < 10 kPa (Fig. 1C, 1D and Fig. S1C).

### Matrix stiffness impacts EV quantity and protein cargo

Above, we determined that tissue stiffness impacts the quantity of vesicles released in breast tumors. Next, we investigated whether matrix stiffness affects EV morphology and protein cargo. Hereafter, we interchangeably refer to EVs released by cells on the plastic matrix as “plastic EVs,” 25 kPa matrix as “stiff EVs”, and 0.5 kPa matrix as “soft EVs”. We compared plastic, stiff and soft EVs derived from highly metastatic, triple-negative-breast-cancer (TNBC) cell lines MDA-MB-231 as our primary model systems (Fig. 2A). We first verified that breast cancer cells displayed the expected stiffness-dependent morphology^19,60^, including a spindle shape on stiffer matrices and a round morphology on the soft matrix, prior to vesicle collection (Fig. S2A).

**Figure 2:**
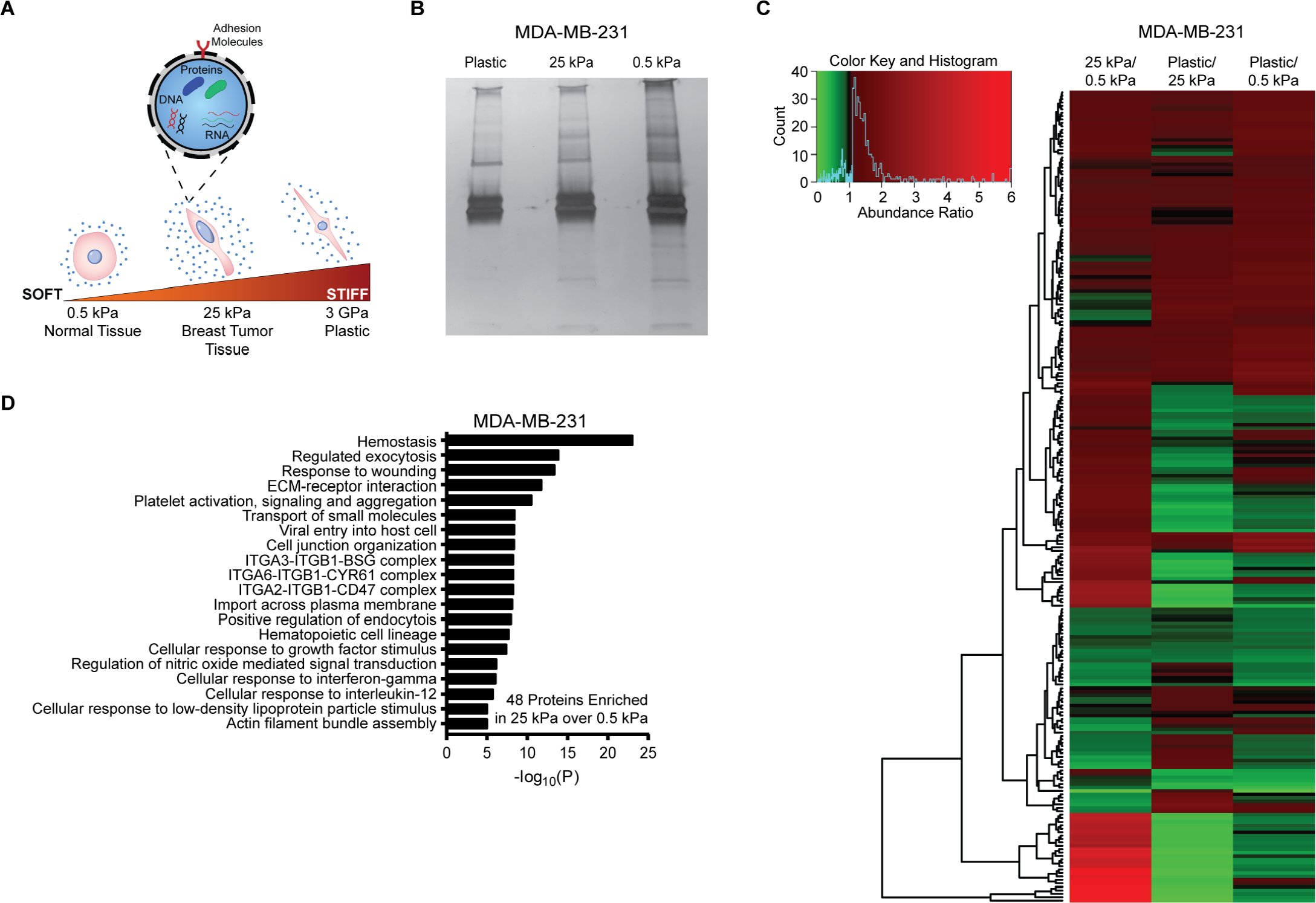
EV cargo is affected by matrix stiffness. **A**, Schematic of EV secretion by cancer cells on standard plastic, tumor (25 kPa) stiffness, and normal-tissue (0.5 kPa) stiffness. **B**, Silver-stain of EV isolated proteins. **C**, Heatmap of EV protein abundance-ratios. Red indicates greater abundance for the condition in the numerator; green indicates greater abundance for the condition in the denominator. Three biological repeats. See also Table S1. **D**, Gene ontology pathway analysis using Metascape for MDA-MB-231 EV proteins enriched in stiff EVs over soft EVs^70^. Three biological repeats.

The size of plastic, stiff and soft EVs was determined using both nanoparticle tracking analysis (NTA) and transmission electron microscopy (TEM). NTA showed that the mean size of collected particles was between 100-150 nm for the TNBC and pancreatic cancer cells across tested matrix stiffnesses (Fig. S2B). Corroborating NTA, TEM indicated that plastic, stiff and soft EVs showed the expected size and morphology of EVs (Fig. S2D)^61,62^. Analysis of the TEM images using a machine learning algorithm^63^ confirmed that the size and shape of EVs were independent of matrix stiffness (Fig. S2E and S2F). Western blots of EV-specific markers^37,64^ confirmed that plastic, stiff and soft EVs contained tetraspanin cluster of differentiation 63 (CD63) and tumor susceptibility gene 101 (TSG101) across all tested matrix stiffnesses (Fig. S2C).

While there were similarities in size and morphology, we found major differences in the protein content of EVs produced by cancer cells on plastic, stiff and soft matrices. We determined that there was a non-significant difference between the total number of vesicles and the amount of isolated protein between EV conditions (Fig. S2G). We then loaded silver-stained electrophoresis gels based on total vesicular protein to normalize for changes in vesicle number and identified qualitative protein cargo differences in the EVs as a function of overall matrix stiffness (Fig. 2B). To test the generality of these findings, we selected pancreatic cancer cell line BxPC3 given that an increase in stiffness is also linked to a poor prognosis in this disease. Pancreatic tumor progression is often characterized by significant changes in the ECM due to desmoplasia, and the selected physiological stiffnesses also match that of normal tissue and extremely stiff tissue in the pancreas^65–69^. We find that the pancreatic cancer cells display the expected morphology on matrices of different stiffness (Fig. S2A). The vesicle size and size distribution are independent of matrix stiffness (Fig. S2B), the vesicles contain CD63 and TSG101 across all conditions (Fig. S2C), and proteins are differentially enriched in the BxPC3 EVs as a function of overall matrix stiffness (Fig. S2H).

To quantify the observed variations in protein content in EVs derived from breast cancer, we performed mass spectrometry on plastic, stiff and soft EVs (Fig. 2C and Table S1). Proteomic analysis of the EVs identified over 200 proteins and revealed significant variations in content between the three conditions (Fig. 2C and Table S1). Using an abundance ratio > 2, we found only 3 proteins enriched in plastic derived EVs over stiff EVs, while 55 proteins were enriched in stiff EVs over plastic EVs (Fig. S3A). When comparing the physiologically relevant stiffnesses, 6 proteins were enriched in soft EVs over stiff EVs, and 48 proteins were enriched in stiff EVs over soft EVs (Fig. S3B). Gene ontology analysis of proteins enriched in stiff EVs identified pathways related to the immune response, tumorigenesis, adhesion, and metastasis including response to wounding, ECM-receptor interaction, cell-junction organization, integrin complexes, and cellular response to interferon-gamma and interleukin-12 (Fig. 2D and Fig. S3D)^70^. In the final comparison, 6 proteins were enriched in plastic EVs relative to soft EVs, and 9 were enriched in soft EVs relative to plastic EVs (Fig. S3C).

The vesicle protein content indicates that plastic, stiff and soft EVs may have different functional roles in promoting metastasis. Given that many of the pathways enhanced in the stiff over soft EVs were related to cell adhesion and cell-ECM interactions (Fig. 2D), we hypothesized that these matrix-stiffness-dependent variations could impact EV biodistribution to different organs (lung, liver, etc.) and the ensuing spread of cancer cells from the primary tumor to these organs. Furthermore, we decided to focus herein on the stiff and soft EVs as the plastic EVs are less physiologically relevant, yielding different cargo that could impact functional assays.

### Stiff EVs show enhanced biodistribution in vivo

To determine whether overall differences in molecular cargo, prompted by differing matrix stiffness, had a functional effect on the distribution and retention of breast cancer derived EVs *in vivo*, we injected immunodeficient nude mice with fluorescent EVs via their tail veins (Fig. 3A). We chose a tail vein injection, which primarily metastasizes to the lung, given our focus on breast cancer: 24 h post intravenous injection of fluorescent EVs, mice were imaged with near-infrared (NIR) from the dorsal, left, ventral, and right sides (Fig. 3B). For all angles, the mean signal-to-noise ratio (SNR) was 2 to 3-fold greater for stiff EVs compared to soft EVs (Fig. 3C). In the lungs, liver, and spleen we observed a 3-fold increase in the mean SNR for stiff EVs over soft EVs (Fig. 3D and 3E).

**Figure 3:**
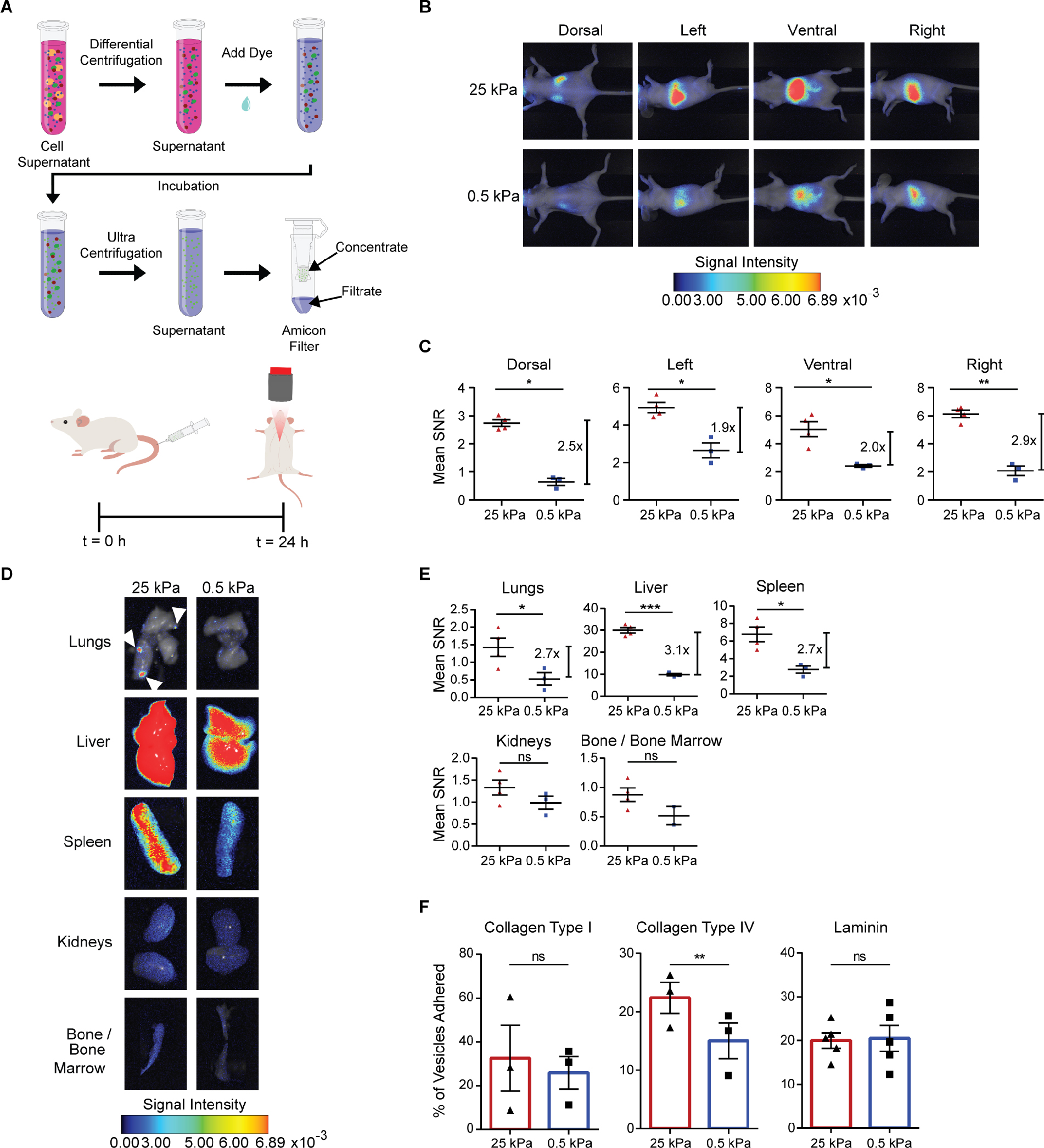
Stiff EVs show increased biodistribution and retention *in vivo*. **A**, Schematic of EV isolation and staining prior to tail-vein injection of 10 μg of EVs in nude mice. **B**, Near-infrared (NIR) imaging and **C**, mean-signal-to-noise ratio (SNR) of MDA-MB-231 vesicle biodistribution in dorsal, left, ventral and right sides (mean ± SEM). Signal intensity is in arbitrary units (a.u.). Three mice in 0.5 kPa condition, and four in 25 kPa condition; one-way ANOVA. **D**, NIR imaging, and **E**, mean SNR biodistribution in the lungs, liver, spleen, kidneys, and bone/bone marrow (mean ± SEM). Signal intensity is in arbitrary units (a.u.). Three mice in MDA-MB-231 0.5 kPa condition and four in 25 kPa condition; one-way ANOVA. **F**, MDA-MB-231 EV binding assay to ECM proteins collagen type I, collagen type IV, and laminin (mean ± SEM). Three biological repeats for collagen type I and collagen type IV; Five for laminin. Paired t-test.

To identify the mechanism driving this stiffness-mediated EV biodistribution, we investigated whether the stiff and soft EVs bound differentially to ECM proteins, especially ECM molecules associated with tumor progression and metastasis^71,72^. Via quantification of total fluorescent signal, stiff EVs preferentially bound to collagen type IV relative to soft EVs (Fig. 3F). The stiff and soft EVs did not demonstrate significant differences in binding to collagen type I or laminin (Fig. 3F). Based on the enrichment of adhesion molecules in stiff EVs in the proteomics data, ITGα_2_β_1_ and CD44 could facilitate the enhanced binding to collagen type IV^72–78^.

### Stiff EVs promote cancer cell dissemination and survival in vivo

Since stiff EVs derived from breast cancer were retained within common secondary sites to a much greater extent than soft EVs, and the stiff EVs bound preferentially to ECM proteins linked to metastasis, we sought to determine whether the EVs would directly affect cancer cell behavior during metastasis. Migration of cancer cells, through directed persistent movement (chemoattraction) and targeted multidirectional movement (chemotaxis), is pivotal for cancer cells leaving the primary tumor site, arriving at the secondary site, and colonizing new tissues^79,80^. Furthermore, EV secretion has previously been linked to cell movement, invasion, and the generation of a pro-tumorigenic and metastatic environment^50,81–86^. We decided to test the chemotactic properties of the breast cancer-derived stiff and soft EVs *in vitro* by using a transwell model (Fig. 4A). Cells were placed on a type I collagen-coated 8 μm pore poly-carbonate membrane with or without EVs in the chamber below (Fig. 4A). After 16 h, 3-times the number of cells migrated towards the stiff EVs than the soft EVs (Fig. 4B and 4C).

**Figure 4:**
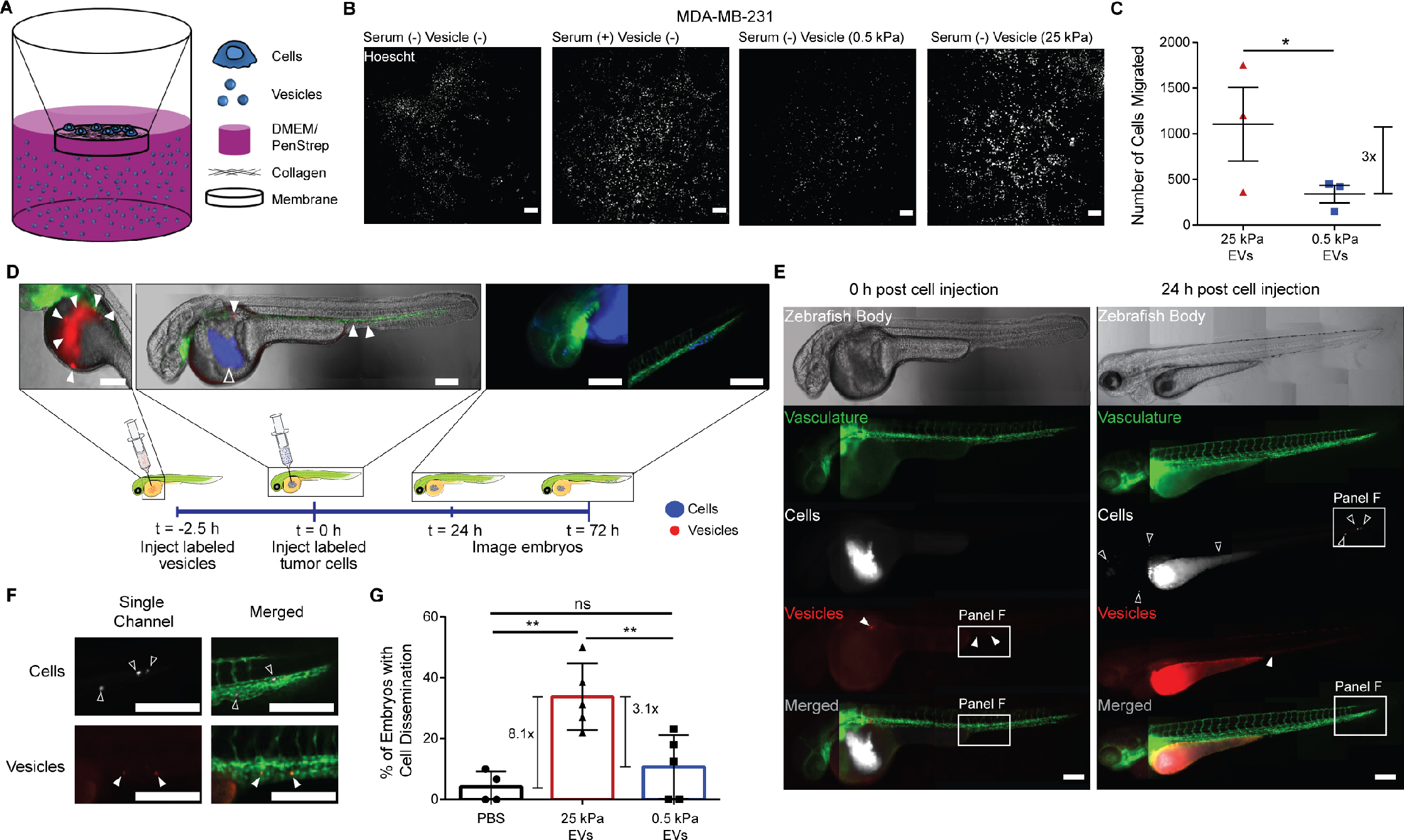
Stiff EVs promote cancer cell migration and dissemination. **A**, Schematic of transwell assay used to test the ability of EVs produced by cancer cells on matrices of different stiffness to differentially promote the recruitment of cancer cells. **B**, Labeled MDA-MB-231 cells that successfully migrated towards the MDA-MB-231 stiff and soft EVs. Scale bar, 100 μm. Representative images. Three biological repeats. **C**, Number of cells migrated to stiff or soft EVs. Three biological repeats. Ratio paired t-test. **D**, Schematic of two-day old zebrafish embryo model expressing green fluorescent vasculature (Tg(fli:EGFP)) used to test the ability of MDA-MB-231 EVs (red, filled triangles) to disperse MDA-MB-231 cancer cells (blue, open triangles). Scale bar, 100 μm. **E**, Full zebrafish body images of the embryos at time 0 h and 24 h post-injection of MDA-MB-231 cancer cells. Scale bar, 100 μm. Representative images. **F**, Enlarged images of disseminated MDA-MB-231 cells (white) and EVs (red). Scale bar, 100 μm. Representative images. **G**, Percentage of injected embryos with cancer-cell dissemination to the head or the tail. The total number of fish per condition is 47 for PBS control, 63 for 25 kPa, and 78 for 0.5 kPa condition. Five biological repeats of EVs. One-way ANOVA.

Enhanced biodistribution (Fig. 3) and chemotactic properties of stiff EVs (Fig. 4C) together would predict a corresponding increase in cancer-cell dissemination *in vivo*. To validate this scenario, we created a zebrafish xenograft model to explore cancer-cell dissemination, survival, extravasation, and migration. Zebrafish possess orthologs for 70% of human genes, are translucent allowing for real-time *in vivo* visualization, cost-effective, and lack adaptive immune systems during early embryogenesis, highlighting their utility as effective xenograft hosts. We utilize zebrafish embryos two days post-fertilization (2 dpf; Fig. 4D).

PBS, breast cancer-derived stiff EVs, or breast cancer-derived soft EVs were injected into the yolk-sac of zebrafish embryos 2 dpf, followed by injection of cancer cells within 2-2.5 h of EV injection (Fig. 4D). Embryos were imaged at 24 h post-cell injection to quantify cell dissemination, and 72 h to observe extravasation (Fig. 4E, 4F, and S4A). Quantification of embryos displaying cancer-cell dissemination to the head and tail 24 h post cell injection in a dye-only condition (no pre-injected vesicles) showed that only 4.2% of fish exhibited a net migration of cells out of the yolk sac to the head or tail of the fish (Fig. 4G). In contrast, 33.8% and 10.7% of fish pre-injected with stiff and soft EVs, respectively, had disseminated cancer cells (Fig. 4G). We also observed that on average 20% of embryos with disseminated cells also had cells that extravasated when treated with 25 kPa EVs and 0% with 0.5 kPa EVs (Fig. S4B). These results show that the role of EVs in mediating metastasis depends critically on the physical properties of the microenvironment.

### Soft EVs transform fibroblasts into CAF-like cells

Since stiff EVs derived from breast cancer facilitate the intravasation, dissemination, and extravasation of cancer cells (Fig. 4), we wanted to assess whether stiff and soft EVs would differentially affect the ability of cancer cells to form tumors at secondary sites by transforming resident stromal cells, particularly fibroblasts. Fibroblasts are responsible for maintaining homeostasis as immunoregulatory cells and through the generation of structural ECM molecules like collagen I^87–89^. Since we observed the greatest differences in EV retention in the lungs (Fig. 3E) and breast cancer frequently metastasizes to the lung *in vivo*, we assessed EV-mediated changes in the phenotype of normal lung fibroblasts (Fig. 5A and 5B). Cancer cells recruited to the lungs are exposed to a relatively soft microenvironment (0.5-1kPa) at this distant site, which has a stiffness similar to that of normal breast tissues^90,91^. To determine how the mechanically new environment of the lung may promote further tumor progression, we investigated how stiff and soft EVs differentially modulated resident lung fibroblasts by assessing the expression of a number of cancer-associated fibroblast (CAF) markers^28,87–89,92–96^. Lung fibroblasts treated with soft EVs experienced a 3.6-fold increase in the expression of CAF-associated molecule^28,87,92–95^ smooth muscle actin (*ACTA2*), 8-fold increase in collagen type I (*COL1A1*), and 3.9-fold increase in vascular endothelial growth factor A (*VEGFA*) compared to those cells treated with stiff EVs (Fig. 5A). Additionally, the fibroblasts exposed to soft EVs showed a 4.6-fold increase in the expression of connective tissue growth factor (*CCN2*), 5.4-fold increase in interleukin 6 (*IL6*), 4.6-fold increase in keratinocyte growth factor (*KGF*), and 4.9-fold increase in vimentin (*VIM*) compared to fibroblasts treated with the stiff EVs (Fig. 5A). While there was a 50% increase in matrix metalloproteinase 1 (*MMP1*) in fibroblasts exposed to soft EVs compared to stiff EVs, this difference was not significant (Fig. 5A).

**Figure 5:**
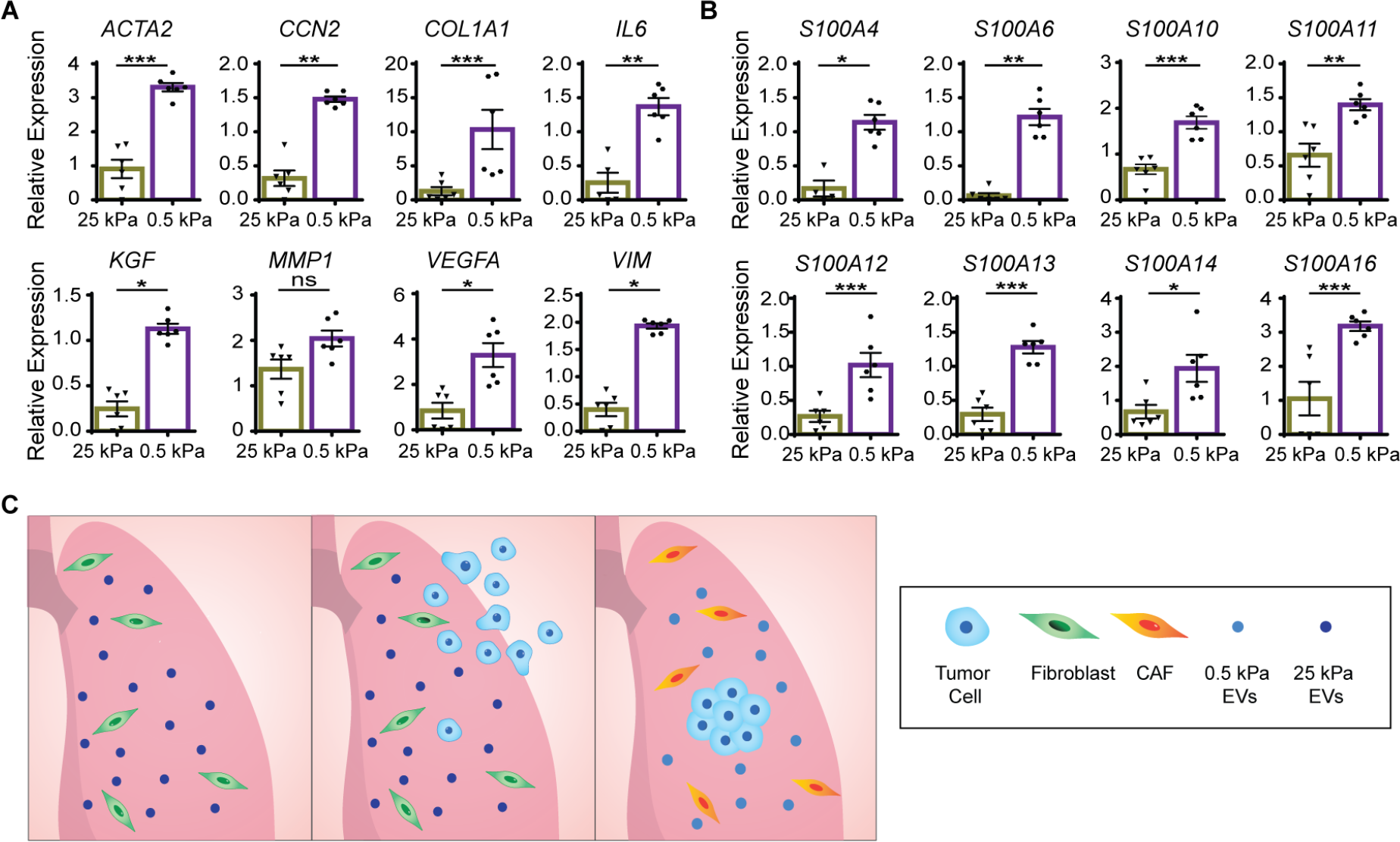
Soft EVs transform the phenotype of resident lung fibroblasts. **A** and **B**, Gene expression fold change in *ACTA2*, *CCN2*, *COL1A1*, *IL6*, *KGF*, *MMP1*, *VIM*, *VEGFA*, and *S100* proteins assessed by qRT-PCR in IMR90 human lung fibroblasts exposed to PBS only, 25 kPa EVs, and 0.5 kPa EVs. Two biological repeats. One-way ANOVA. **C**, Schematic showing (left) the arrival of stiff EVs in the lung, a mechanically soft environment, and encountering resident normal lung fibroblasts; (middle) cancer cells are then recruited to the lung; and (right) the cells, now experiencing a soft environment, release soft EVs that transform the resident fibroblasts to a cancer-associated fibroblast (CAF) phenotype.

We then probed how stiff and soft EVs regulated normal lung fibroblast expression of S100 proteins. In breast and pancreatic cancers, dysregulation of S100 protein expression, due in part to CAFs, is tied to an increase in growth, metastasis, and angiogenesis^97^. Previously, the success of pre-metastatic niche formation in the lung was determined to be dependent on S100 protein upregulation^98^. In lung fibroblasts exposed to soft EVs, we observed a noticeable up-regulation of four S100 genes (*S100A10*, *S100A11*, *S100A14*, *S100A16*) (Fig. 5B). Our results also indicate that stiff EVs downregulate the expression of *S100A4*, *S100A6*, *S100A12*, and *S100A13* resulting in a respective 6.8-fold, 19.2-fold, 3.8-fold, and 4.3-fold difference relative to soft EVs (Fig. 5B). Together, these results suggest that soft vesicles produced by newly disseminated cancer cells in the soft microenvironment of the lung are significantly more effective at producing a CAF-like phenotype in lung fibroblasts via ensuing up-regulation of S100 inflammatory signals and *ACTA2*/*COL1A1*/*VEGFA* (Fig. 5C).

## Discussion

This work demonstrates the importance of utilizing physiologically relevant conditions for studying the role of EVs in cancer. EVs released by cancer cells on plastic dishes differentially expressed hundreds of proteins, resulting in inaccurate information about the ability of vesicles to distribute *in vivo* and promote cell dissemination compared to matrices that mimic tissue stiffness at the primary and distant sites (Fig. S3D and S5). Our stiff tumor tissue and soft normal tissue matrices significantly altered EV quantity, protein cargo, function, and their potential to affect multiple aspects of the metastatic cascade.

The quantification of vesicles from primary patient breast tissue indicates that more EVs are released in stiff tissue over soft tissue. This increased EV release is consistent with stiffness mediated Rab8 activation promoting vesicle biogenesis^99^. Within the tissue, there are many different cell types, all contributing to the number of small EVs we isolated in this study. As of now, there are no effective methods or markers for separating EVs based on the cell type of origin, which limits our ability to determine the number of vesicles produced by each cell type of the tumor microenvironment. We do notice though that there is an increase in cell number and density within the stiff breast tumor tissue compared to the soft breast tumor tissue^20^.

In addition to the observed variations in stiffness-dependent vesicle secretion, the protein cargo of EVs critically depends on matrix stiffness. We identified a 3-fold increase in mean SNR between the stiff and soft EVs in the lungs and liver, two of the most common sites of breast cancer metastasis. These results suggest that small EVs have a different rate of clearance *in vivo* as a function of stiffness, presumably due to our observed stiffness-dependent presentation of adhesion molecules on EVs and their ECM binding affinity (Fig. 6A). The increased retention of stiff EVs in the lung could be a function of specific integrins, including α_6_β_4_ and α_6_β_1_, which have been previously linked to organotropic homing^49^. Collagen type I, collagen type IV, and laminin have all been linked previously to cancer cell migration and invasion^71–73,100–106^. Collagen type IV lines all basement membranes in the liver and airway basement membrane in the lung^107,108^. Preferential binding of stiff EVs to basement-membrane proteins may, therefore, also explain increased stiff EV retention in the lungs and liver. Previously, small EVs were shown to increase lung vascular permeability *in vivo*^49^. Our results suggest that stiff EVs, which bind these basement-membrane proteins and promote chemotaxis, draw cancer cells to the lung.

**Figure 6:**
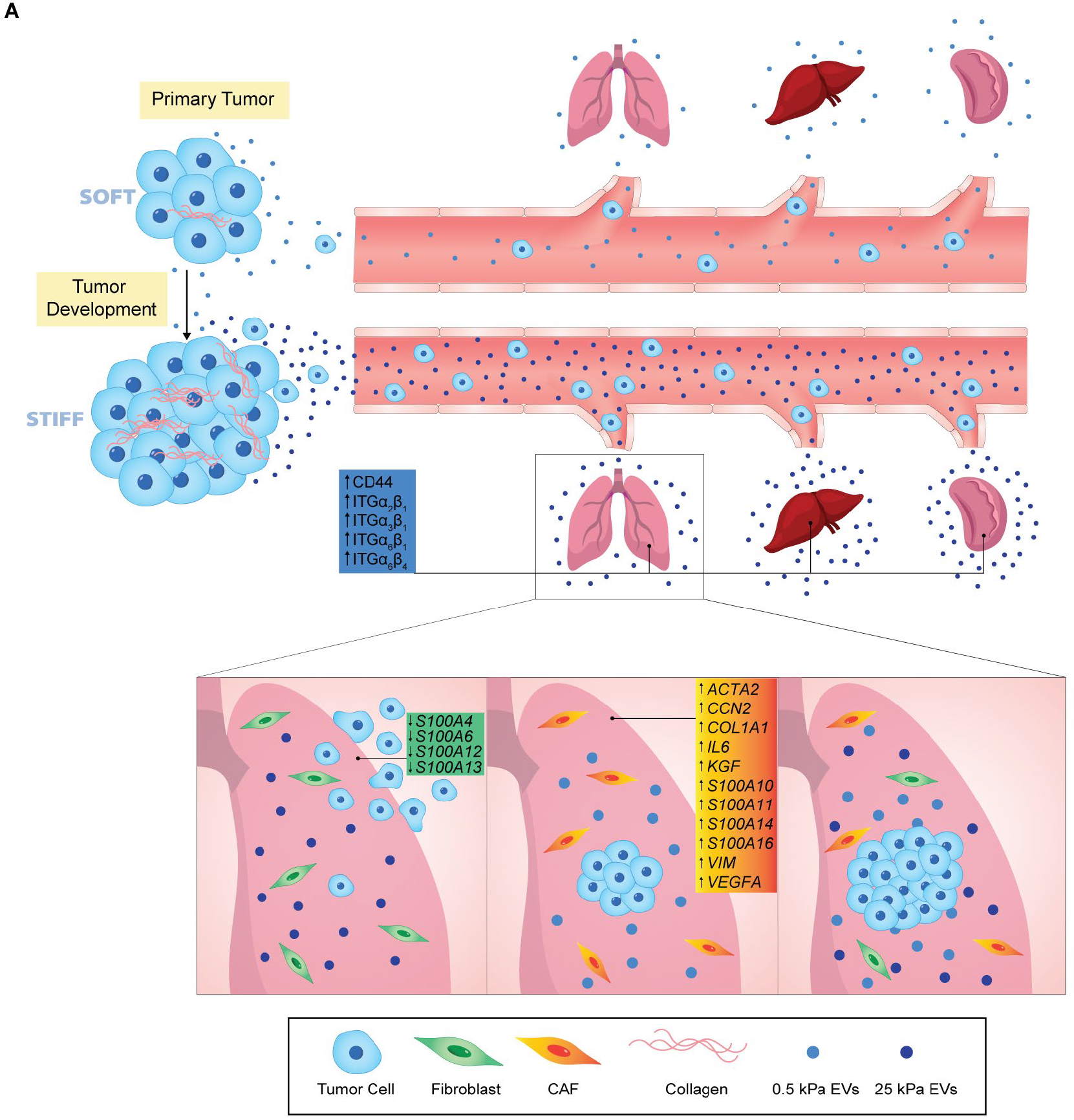
Matrix stiffness alters EV protein cargo and affects EV functions in metastasis. **A**, Schematic describing the overall impact of stiff and soft EVs on the formation of metastasis. As the primary tumor develops, cancer cells (blue cells) experience a change in matrix stiffness, from soft to stiff. Initially, in a soft environment, cancer cells release soft EVs (purple) that create a pro-tumorigenic environment. Now in a stiff matrix environment, cancer cells release stiff EVs (blue) that have an enhanced ability to distribute and reside (blue text boxes) in distant organs (i.e., lungs, liver, kidneys). These stiff EVs can promote cancer cell migration and dissemination (blue text boxes). Resident normal lung fibroblasts (green cells), upon taking up stiff EVs, decrease their expression of pro-inflammatory markers *S100A4*, *S100A6*, *S100A12*, and *S100A13* (green text box). Once cancer cells have metastasized to the lung, they re-experience a soft matrix and release soft EVs that transform the resident fibroblasts into a CAF phenotype (yellow/orange cells) by increasing expression of *ACTA2*, *COL1A1*, *VEGFA*, and several other genes (yellow/orange text box). Inducing this fibroblast phenotype allows for the cancer cells to colonize and proliferate in a pro-tumorigenic environment.

Based on transwell and zebrafish experiments, the mechanism behind EV-mediated cell movement depends on matrix stiffness. Compared to soft EVs, stiff EVs demonstrated an enhanced ability to induce cancer cell migration *in vitro* and dissemination *in vivo* (Fig. 4C and 4G). Furthermore, we noticed that cancer cells pre-treated with stiff EVs seemed to extravasate and migrate into the zebrafish tissue to a greater extent than cells pre-treated with plastic or soft EVs (Fig. S4B). These findings suggest that the proteins responsible for cell spreading from the primary tumor are preferentially sorted into EVs released by cancer cells experiencing a stiff tissue matrix.

While stiff EVs are more effective at promoting the early and middle steps of the metastatic cascade through migration, dissemination, and arrival at distant tissues, we determined that stiff and soft EVs operate in a dynamic way to colonize distant organs, especially the lung. Once internalized by normal lung fibroblasts, the stiff EVs downregulate S100 expression, while soft EVs upregulate activation, vasculogenic, and inflammatory markers in the fibroblasts. The increased retention of stiff EVs in the lung and the downregulation of S100 proteins in normal resident lung fibroblasts may seem counterintuitive; however, a decrease in S100A4 expression has been linked to blocking fibroblast invasion and T-cell recruitment at the primary tumor^109^. Additionally, there is a direct relationship between the expressions of VEGFA and S100A4 in fibroblasts, with the expression of both molecules being important for metastatic colonization^110^. Decreased *S100A6* expression in breast cancer has been linked to a worse prognosis regardless of subtype^111^. Although little has been studied about its role in breast cancer^111^, S100A13 is known to regulate fibroblast growth factor (FGF1) and interleukin 1α (IL1α), which can affect the angiogenic and mitogenic properties of the tumor microenvironment^112–114^. Therefore, the decrease in the expression of these S100 proteins in fibroblasts can promote a pro-metastatic environment in the lung (Extended Data Fig. 6a), prior to the arrival of cancer cells.

Once cancer cells arrive in the new soft environment of the lung, they release soft EVs that transform the resident fibroblasts towards a CAF phenotype through increased expression of *ACTA2*, *COL1A1*, and *VEGFA* (Fig. 6A). This interaction between the soft environment and fibroblasts could also take place during early tumor progression or at other distant sites of metastasis^48,54–59^. The soft EVs demonstrate an upregulation of S100, cytoskeletal regulating, binding, and cell signaling proteins linked to primary tumor growth (Fig. 5A and 5B). We propose that stiff EVs direct the recruitment of cancer cells and ensure retention in the lung by generating an anti-inflammatory environment; once there, the cancer cells experience a soft matrix and release soft EVs that transform the surrounding stroma into a pro-tumorigenic environment (Fig. 6A).

Together our results indicate that EVs promote metastasis through multiple mechanisms that take advantage of the differences in stiffness of the primary and metastatic sites (Fig. 6A). The first is through the increased retention and biodistribution of stiff EVs *in vivo*, due to augmented binding to the ECM via increased integrin presentation, which allows for the formation of pre-metastatic niches. Second, through the generation of chemotactic gradients, stiff EVs promote cancer cell recruitment to metastatic sites *in vivo*. Third, EVs produced at both normal and tumor tissue stiffnesses affect changes to the surrounding ECM by regulating fibroblast activity. Stiff EVs decrease inflammatory signaling in the fibroblasts to facilitate cancer cell arrival at the lung, and once inside the mechanically soft lung, cancer cells release soft EVs that increase the expression of pro-tumorigenic markers in the fibroblasts. Our findings highlight the critical importance of the physical properties of the ECM on the quantity, quality, and function of EVs produced by cancer cells in mediating their metastasis. Future exploration may focus on investigating pan-cancer markers of EV-driven stiffness-mediated metastasis for diagnostic and therapeutic applications.

## Acknowledgements

We would like to thank Dr. Simion Kremer and Dr. Bob Cole at the Mass Spectrometry and Proteomics Facility for the Johns Hopkins University School of Medicine for running the proteomics experiment and writing the methods section. Thank you to Prof. Alan Meeker at the Johns Hopkins Oncology Tissue Services Core for his insight and guidance throughout the project. We thank Dr. Barbara Smith in the Kuo Microscopy Facility for taking TEM images and writing the corresponding methods section. We thank Vishnu Prasath, Dr. David Wilson, Prof. Jordan Green, Prof. Karen Reddy, Lindsay Rizzardi, Kimberly Stephens, the Sean Taverna Lab, the Andrew Feinberg Lab, Dr. Rada Cordero-Gonzalez, Prof. Arturo Casadevall, Dr. Josh DiGiacomo, and Prof. Daniele Gilkes for their expert help and access to resources for the project. We also thank Giulianna Leotta, Dr. Michael Harris, Prof. Ashley Kiemen, Dr. Meng Horn Lee, Prof. Pei-Hsun Wu, Adrian Johnston, Bailey Robertson, Prof. Jerry S.H. Lee (University of Southern California), Prof. Ken Witwer (Johns Hopkins University), and Prof. Martin Humphries (The University of Manchester) for fruitful discussions.

This work was supported by grants from the National Cancer Institute (U54CA210173, U54CA143868, U54CA268083, R01CA174388), the National Institute on Aging (U01AG060903) and the National Institute of Arthritis & Musculoskeletal & Skin Diseases (U54AR081774) to D.W.; National Science Foundation Graduate Research Fellowship (1746891) to A.S. Additionally, the NVIDIA Corporation with the donation of the Titan X (Pascal) GPU (to AMB), Ministerio de Ciencia, Innovación y Universidades, Agencia Estatal de Investigación, under grants TEC2015-73064-EXP, TEC201678052-R and PID2019-109820RB-I00, MINECO/FEDER, UE, co-financed by European Regional Development Fund (ERDF), “A way of making Europe” (to AMB), and a 2017 Leonardo Grant for Researchers and Cultural Creators, BBVA Foundation (to AMB).

## Author Contributions

A.S. formulated the hypothesis, designed, and performed experiments, analyzed, and interpreted the data, managed the project, and wrote the manuscript. Y.L., G.E.C., and T.S.K.E. performed the zebrafish experiments. C.M., N.F, D.M.L., S.K., S.I., M.I., H.H., B.S., W.D., R.V., G.R., M.K. performed experiments. B.S., G.R., M.K., and P.R. performed the mouse experiments. J.K. and L.G. performed mechanical testing on primary patient tissue samples. A.L. designed and created the schematics in the paper. S.K., D.M.L., and M.I. contributed to formatting the figures. E.G.M. and A.M.B. performed the computational analysis of the EVs in TEM images. M.H. provided primary patient tissue. L.G. and T.S.K.E. contributed to the experimental design and interpretation of the data. D.W. formulated the hypothesis, managed the project, interpreted the data, and wrote the manuscript.

## Materials and Methods

### Cell culture for EV collection

Human breast cancer cell lines MDA-MB-231, SUM149, and MDA-MB-468, human pancreatic cancer cell line BxPC3 (all from ATCC), and IMR90 human lung fibroblasts (gift from Daniele Gilkes, Johns Hopkins University) were cultured in 013-CV DMEM (Corning) containing 10% fetal bovine serum (Corning) and 1% penicillin-streptomycin (Gibco). After a 24 h incubation at 37°C, cells were washed with Dulbecco’s phosphate buffered saline (DPBS, Corning) and changed to 013-CV DMEM containing 10% exosome-depleted fetal bovine serum (Gibco) and 1% penicillin-streptomycin (Gibco). Prior to vesicle collection, cells were seeded at 70% confluency and cultured on 177 cm^2^ 0.5 kPa Collagen Type I Coated Plates (Matrigen), 177 cm^2^ 25 kPa Collagen Type I Coated Plates (Matrigen), and 150 cm^2^ plastic tissue culture flasks (“plastic”) (Falcon). Following another 24 h incubation at 37°C, EVs were collected and purified.

### EV collection

Cell culture supernatant was subjected to sequential centrifugation (800 × g for 5 min, 2,000 × g for 10 min, 10,000 × g for 30 min) at 4°C (Beckman Coulter Avanti J-E Centrifuge); filtered through a 0.22 μm PES filter (Genesee); centrifuged twice at 100,000 × g for 2 h at 4°C (Beckman Coulter Optima XE-90 Ultracentrifuge). The supernatant was replaced with DPBS between ultracentrifugation spins. The final vesicle pellet was re-suspended in 1 mL DPBS. Samples were concentrated using a 2 mL 3 kDa Amicon filter (MilliporeSigma). For the mouse experiments, samples were incubated with 1 μM DiR dye (Thermo Fisher) prior to ultracentrifugation. For the zebrafish studies, EVs were labeled with the CMTPX Dye (Thermo Fisher Scientific) prior to concentration using an Amicon filter. Protein concentration was determined using a Pierce BCA protein assay kit (Thermo Fisher Scientific) according to the manufacturer’s protocol.

### Size distributions of EVs

Size distribution and concentration of EVs were measured using a NanoSight NS300 (Malvern Preanalytical). Additional details are in supplementary information.

### Western blot

EV protein aliquots were lysed by boiling with 20% β-mercaptoethanol in 4X Laemmli buffer (Bio-Rad) for 5 min at 100°C. Lysates were separated by molecular weight on 4-15% Mini-Protean Precast TGX Gels (Bio-Rad) and transferred to Trans-Blot Turbo Mini PVDF membranes (Bio-Rad). Overnight incubation at 4°C with primary antibodies, including anti-TSG101 (1:500, ProteinTech) and anti-CD63 (1:1000, Abcam) in 1X TBST. Secondary antibody incubation with corresponding HRP-conjugated anti-rabbit or anti-mouse secondary antibody (Cell Signaling Technology).

### Silver stain

Samples were prepared according to the western blot protocol and run through a 4-15% Mini-Protean Precast TGX Gel (Bio-Rad). The gels were then stained using the Pierce Silver Stain Kit (Thermo Fisher Scientific) according to the manufacturer’s protocol.

### Transmission electron microscopy

10 µL of sample was adsorbed to glow-discharged (EMS GloQube) 400 mesh ultra-thin carbon coated grids (EMS CF400-CU-UL) for 2 min, followed by three quick rinses of TBS and stained with 1% UAT (uranyl acetate with 0.05% Tylose). Grids were immediately observed with a Philips CM120 at 80 kV and images captured with an AMT XR80 high-resolution (16-bit) 8 Mpixel camera. Two biological repeats.

### Biodistribution of EVs in mice

All mouse work was performed following Johns Hopkins University and IACUC guidelines under animal protocol MO16A383. There was no statistical method to pre-determine sample size. 6-8-week-old NCr nude (NCRNU-F sp/sp) females (Taconic) were each injected via tail vein with stained vesicles in DPBS at a quantity of 10 μg of protein in 50 μL per mouse. 24 h post injection, mice and their organs were imaged using the LI-COR Pearl Impulse Imaging System (LI-COR Biosciences). Images were analyzed in LI-COR Pearl Impulse Imaging System according to the manufacturer’s instructions. The mean signal to noise ratio (SNR) was determined by subtracting the mean background intensity from the mean intensity of the region of interest and dividing through by the standard deviation of the background.

### Biodistribution of EVs and cancer cells in zebrafish

All procedures on zebrafish (Danio rerio) were approved by IACUC at The University of Pennsylvania. Fertilized zebrafish eggs of the transgenic strain expressing enhanced green fluorescent protein (EGFP) under the fli promoter (fli:EGFP) or mCherry under the flk promoter (flk:mCherry) were incubated at 28 °C in E3 solution and raised using standard methods. Embryos were transferred to E3 solution containing 5 µg/ml proteases and 0.2 mM1-phenyl-2-thio-urea (Sigma) 24 h post-fertilization to dechorionate the fish embryos and prevent pigmentation, respectively. At 48 h post-fertilization, zebrafish embryos were anesthetized with 0.03% tricaine (Sigma) and then transferred to an injection plate made with 1.5% agarose gel for microinjection. Approximately 200,000 EVs suspended in PBS were injected into the yolk sac of each embryo using a XenoWorks Digital Microinjector (Sutter Instrument). Each injection volume was between 5-10 nL. At 2-2.5 h post the vesicle injection, 150 – 400 MDA-MB-231 cells labeled with NucBlue live cell stain ReadyProbe (Thermo Fisher Scientific) and suspended in complete growth medium supplemented with 0.5 mM EDTA were injected into the yolk sac of each vesicle-bearing embryo. Pre-pulled micropipettes were used for the microinjection (Tip ID 50 µm, base OD 1 mm, Fivephoton Biochemicals). After injection, the fish embryos were immediately transferred to a PTU-E3 solution. Injected embryos were kept at 33 °C and were examined every day to monitor tumor migration using a widefield microscope. Images for extravasation analysis were taken using an Olympus spinning disk confocal microscope.

### Patient tissue sample preparation for mechanical measurements

All patient tissue samples were obtained with written consent from the patient and approved by the Johns Hopkins Medicine Institutional Review Board (IRB). Tissue samples received from the patients were kept at 4°C DPBS immediately after mastectomy or lumpectomy. Tumor samples were then transferred for mechanical tests within 4 h of resection. The tumor tissue was then sectioned to expose the regions of interest for micromechanical mapping and compression tests.

### Tumor stiffness mapping using microindentation

Dynamic indentation using a nanoindenter (Nanomechanics, Inc.) was used to characterize the tumor elastic modulus ^115^. Sneddon’s stiffness equation ^116^ was applied to relate the dynamic stiffness of the contact to the elastic storage modulus of the samples ^117,118^. Additional details are in the supplementary information document.

### Compression tests of tumor-adjacent and tumor tissues

Compression tests were performed as previously reported ^20^. Briefly, tissue samples were sectioned to obtain flat and parallel surfaces on all sides. Once the sample was sectioned, it was immediately staged on a tensile/compression tester (MTS Criterion) for measurement ^119^. The top compression plate was lowered until in full contact with the tissue sample at a minimal load. Once in contact, the samples could relax and stabilize for 1 min before the actual compression test. Tissue samples were compressed at 0.25 mm/sec deformation rate until 20% strain. Young’s modulus calculation was done on the best-fitted slope of the initial linear region (∼5-10%) of the obtained stress-strain curve. A single measurement was obtained for each tissue.

### Vesicle collection and characterization for patient tissues

After mechanical measurements, tissue was transferred to 5 mL of 1% penicillin-streptomycin solution in 013-CV DMEM and incubated at 37°C overnight. After 24 h, tissue was fixed in formalin, and vesicles were isolated from the supernatant. Additional details are in the supplementary information document.

### ECM binding assay

Substrates were obtained from Millicoat ECM Screening Kit (MilliporeSigma) and rehydrated according to manufacturer specifications. 1.5×10^9^ CMTPX fluorescently labelled vesicles in 50 μL DPBS (without Ca^2+^ & Mg^2+^) were incubated on each substrate for 1 h at 37°C. Wells were imaged at 10X TRITC channel with 100% laser intensity and 100 ms exposure time (Nikon Eclipse Ti), and fluorescence was determined in ImageJ by measuring the mean intensity of a fixed region of interest. After removing the diluted suspension, the matrix was washed three times using DPBS (with Ca^2+^ & Mg^2+^) according to the manufacturer protocol, and the wells were imaged again under DPBS (without Ca^2+^ & Mg^2+^) to minimize possible fluorescence deviation from ions. For all washing steps, slow manual pipetting was adopted to avoid disturbance of the adhered EV samples. Background intensity was determined from the negative control – PBS with CMTPX dye processed through the same ultracentrifugation and 3kDa Amicon filtration steps as EV samples – and subtracted from sample intensity. Sample intensity post-wash was divided by the pre-wash intensity values at the same ROI to determine the percentage of vesicles adhered to each substrate.

### Fibroblast mRNA expression assay

IMR90 lung fibroblasts were seeded two days prior to the addition of vesicles. These cells were then washed with DPBS and incubated in an exo-depleted medium with vesicles for 48 h at 37°C. RNA was extracted according to manufacturer instructions for DirectZol Kit (Zymo Research) after imaging (Nikon Eclipse Ti). cDNA was generated using iScript cDNA Kit (Bio-Rad) according to manufacturer instructions. Then qPCR was performed. Two biological repeats with three technical repeats per condition. Housekeeping gene value is a geometric mean of α-tubulin (*TUBA3C*), glyceraldehyde-3-phosphate dehydrogenase (*GAPDH*), and TATA-Box Binding Protein (*TBP*). The primer sequences used are listed in Table S2.

### Transwell assay

8 μm pore polycarbonate transwell inserts (Olympus Plastics) were coated with 50 μg/mL collagen type I solution and incubated for 1 h at 37°C. Post-incubation, wells were washed with DPBS, and MDA-MB-231 cells were seeded in the top chamber in a serum-free medium. The plate was then incubated for 1 h at 37°C to allow the cells to settle onto the membrane. Next, various conditions including a serum-free negative control, serum-containing positive control, and stiff (25 kPa) or soft (0.5 kPa) vesicle-containing conditions were added to the lower chamber. After 16 h incubation at 37°C, cells on the bottom side of the membrane were stained with a 1:100 dilution of Hoescht (Thermo Fisher Scientific) and imaged at 4X and 10X (Nikon Eclipse Ti). The number of cells detected was quantified using ImageJ.

### Quantifying EV secretion

To determine quantifiable variations, NTA particle concentrations were multiplied by UC sample volumes for a total particle number. Dividing the total vesicle number by the weight of the tissue provides a value of vesicles secreted per gram of tissue.

### EV proteomics

10-plex TMT was performed on three biological replicates of EVs from cells cultured on tissue culture plastic, 25 kPa, and 0.5 kPa matrices^120^. Data was searched using SwissProt *Homo Sapiens* database with MASCOT in Proteome Discoverer 2.2 (Thermo). Additional details are in the supplementary information document.

### Statistical Analysis

Statistical analysis was performed using Prism 6 (GraphPad Software, Inc.) to calculate the mean, standard deviation, and standard error mean. T-test and One-way ANOVA were performed where appropriate to determine significance (GraphPad). Biological and technical replicates are indicated throughout the figure captions. All graphical data are reported as mean ± SEM. * p < 0.05, **p < 0.01, *** p< 0.001, and *** p< 0.0001.

### Data Availability Statement

The datasets that support the findings of this study are available from the corresponding author upon request.

