## Supplementary material for "Small extracellular vesicles promote stiffness-mediated metastasis": Fig. S

### Supplementary Information

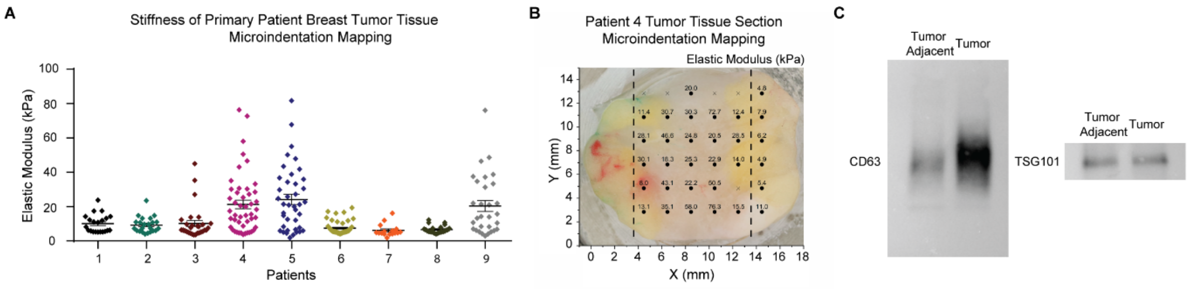


**Supplementary Figure 1: Stiffness measurements of primary patient tumor tissues**

**A**, Microindentation measurements (kPa, mean ± SEM) for each breast-cancer patient tissue tumor sample. Nine patients. **B**, Microindentation mechanical mapping of a primary patient breast tumor tissue sample. Dark circles indicate measurements of the elastic modulus expressed in units of kPa. Crosses are non-measurable indentations. The dotted lines indicate where the tissue was sectioned into stiff (middle) and soft (right) regions for vesicle collection. **C**, Representative western blots of EV markers CD63 and TSG101 for EVs isolated from patient tumor adjacent and tumor tissue.

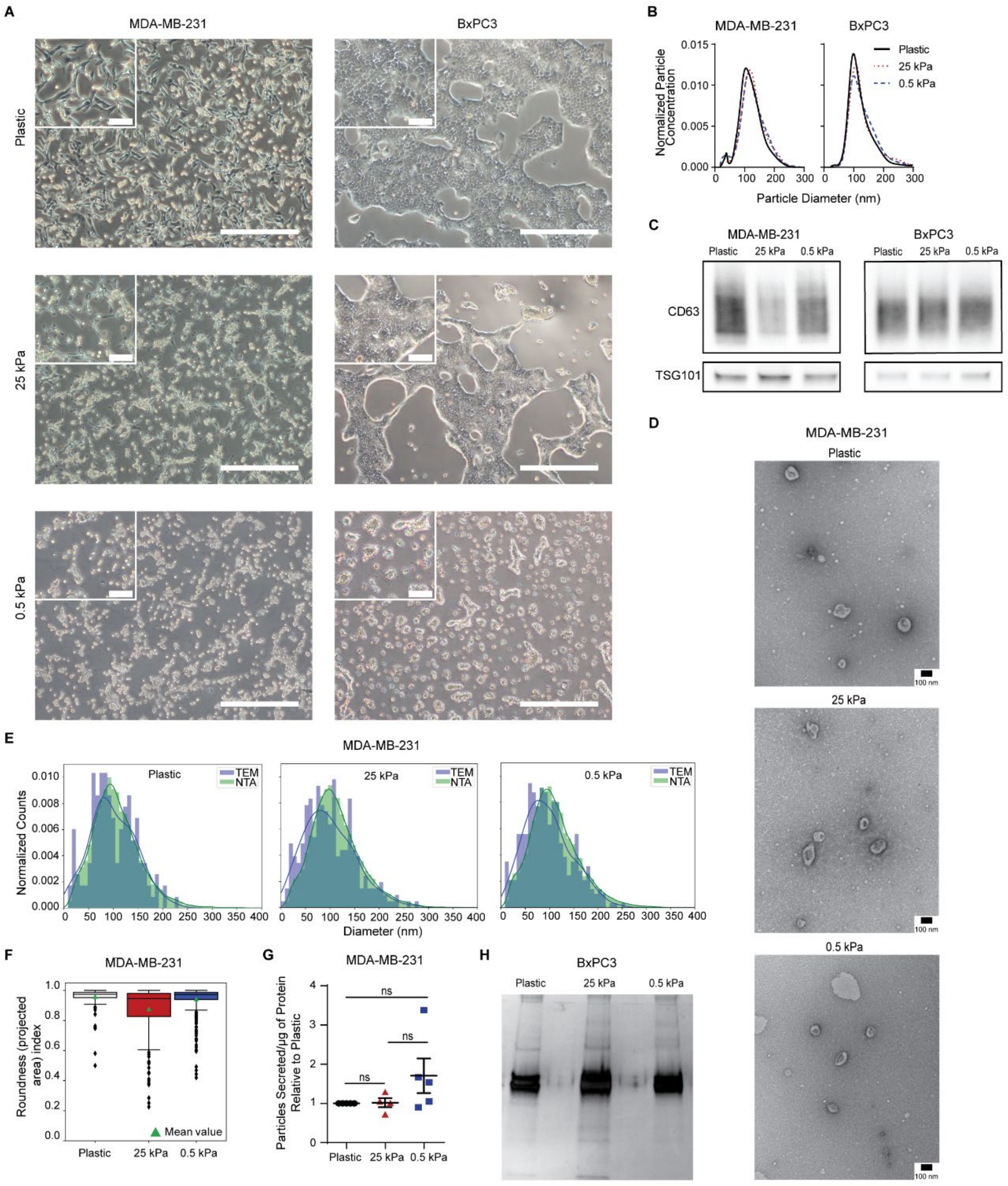


**Supplementary Figure 2: Isolation and characterization of EVs produced by cancer cells *in vitro***

**A**, Morphology of MDA-MB-231 human triple negative breast cancer cells and BxPC3 human pancreatic cancer cells cultured on tissue culture plastic, 25 kPa (stiffness of breast tumor tissue), and 0.5 kPa (stiffness of normal breast tissue) matrices. Scale bar, 500 μm. Inset scale bar, 100 μm. **B**, Size distribution of vesicles released by human triple negative breast cancer cells (MDA-MB-231) and human pancreatic cancer cells (BxPC3) on plastic dishes and on 25 kPa, and 0.5 kPa collagen I-coated matrices. **C**, Representative western blots of EV markers CD63 and TSG101 for EVs produced by MDA-MB-231 and BxPC3 cells on plastic dishes and on 25 kPa and

0.5 kPa matrices. Nine biological repeats. **D**, Representative TEM images of EVs produced by MDA-MB-231 cells. Scale bar, 100 nm. Two biological repeats. **E**, Comparison of the size distribution of vesicles produced by MDA-MB-231 cells measured via NTA and machine-learning-based TEM. Two biological repeats. **F**, Roundness (projected area) index of EVs from TEM images. Two biological repeats. **G**, Number of EVs released per microgram of protein relative to the plastic condition. Five biological repeats plastic EVs, four biological repeats stiff EVs, and five biological repeats soft EVs. **H**, Silver-stain of BxPC3 EV isolated proteins.

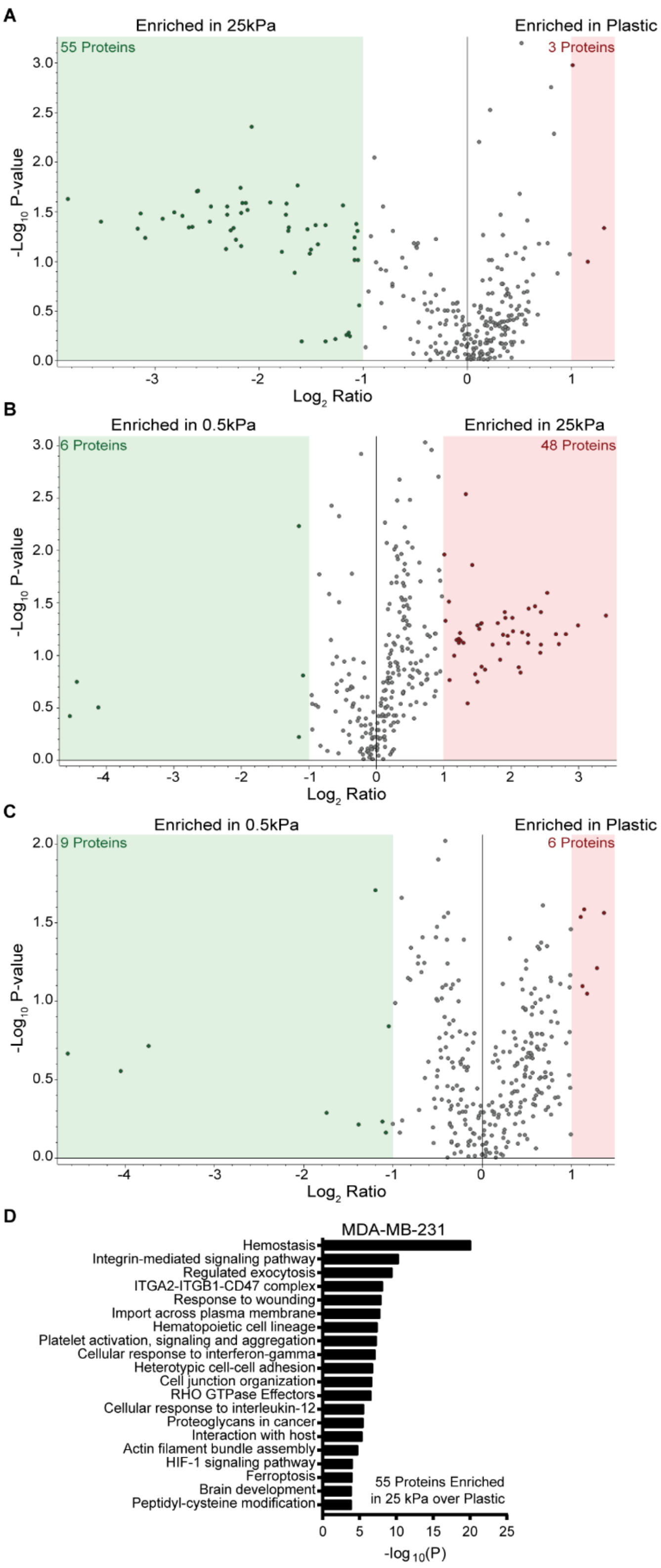


**Supplementary Figure 3: Protein cargo is different for plastic, stiff and soft EVs**

Volcano plots of differentially abundant proteins between (**A**) plastic and 25 kPa EVs, (**B**) 25 kPa and 0.5 kPa EVs, and (**C**) plastic and 0.5 kPa EVs. The x-axis is the $\log_2$ ratio. The y-axis is the $-\log_{10}$(P-value). Red denotes those proteins enriched >2-fold in the vesicle condition in the numerator. Green denotes those proteins enriched >2-fold in the vesicle condition in the denominator. The number of enriched proteins is written in red and green. Three biological repeats. **D**, Gene ontology pathway analysis using Metascape for the 55 proteins enriched in the stiff EVs over the plastic EVs by MDA-MB-231 human breast cancer cells (Zhou *et al.*, 2019). Three biological repeats.

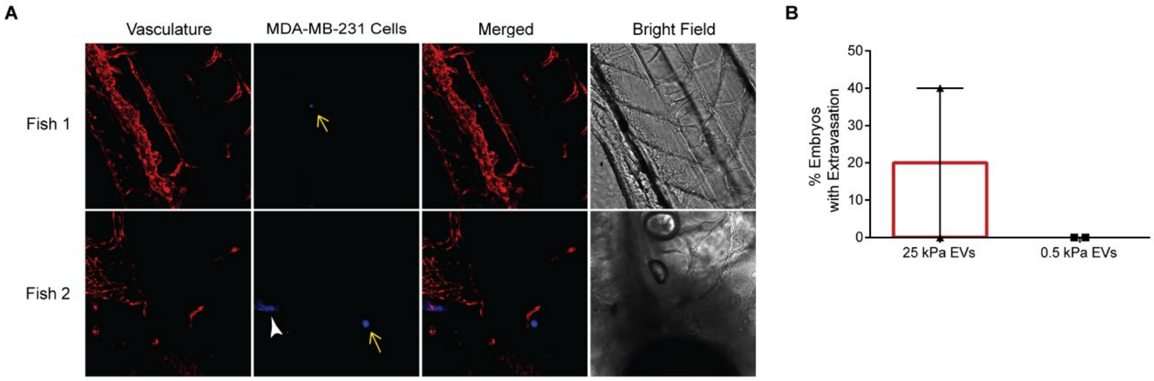


**Supplementary Figure 4: Stiff EVs promote extravasation**

**A**, Images of zebrafish 72 h after cell injection (vasculature = red, MDA-MB-231 cells = blue). Yellow arrows denote single cells. White arrow denotes autofluorescence of a fish structure. **B**, Stiff EVs promote cell extravasation 20% of the time, while soft EVs promote cell extravasation 0%. Two biological repeats of EVs.

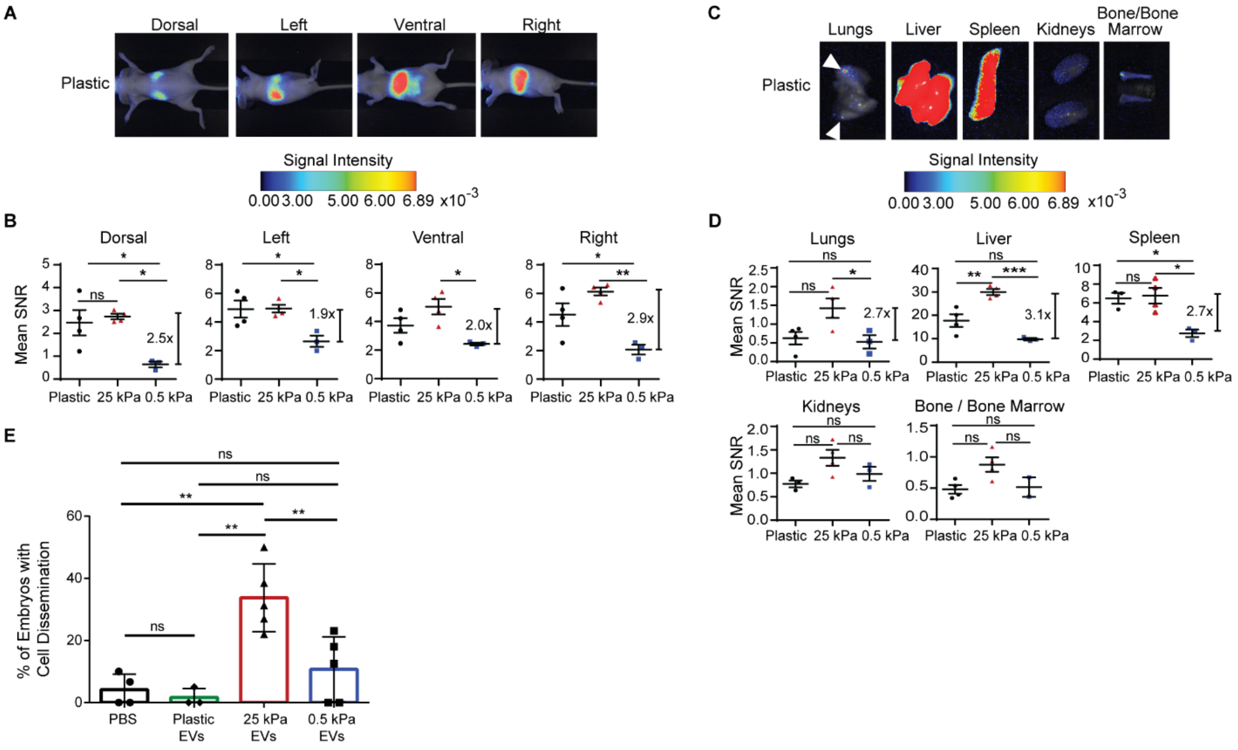


**Supplementary Figure 5: Plastic EVs differ in effect from stiff and soft EVs**

**A**, Near-infrared (NIR) imaging and **B**, mean-signal-to-noise ratio (SNR) of MDA-MB-231 vesicle biodistribution in dorsal, left, ventral and right sides (mean ± SEM). Signal intensity is in arbitrary units (a.u.). Three mice in 0.5 kPa condition, and four in plastic and 25 kPa conditions; one-way ANOVA. **C**, NIR imaging and **D**, mean SNR biodistribution in the lungs, liver, spleen, kidneys, and bone/bone marrow (mean ± SEM). Signal intensity is in arbitrary units (a.u.). Three mice in MDA-MB-231 0.5 kPa condition, and four in plastic and 25 kPa conditions; one-way ANOVA. Only three mice are shown for plastic in the spleen and kidneys plots since one mouse in plastic condition was missing images for the spleen and kidneys. **E**, Percentage of injected embryos with cancer-cell dissemination to the head or the tail. Total number of fish per condition is 47 for PBS control, 40 for plastic, 63 for 25 kPa, and 78 for 0.5 kPa condition. Three biological repeats of EVs for plastic and five for 25 kPa and 0.5 kPa EVs. One-way ANOVA.

**Supplementary Tables**

**Table S1: Figure 1C heatmap dendrogram row labels in order from top to bottom**

| | Gene Symbol | | Gene Symbol | | Gene Symbol |
|---|---|---|---|---|---|
| 1 | CP | 80 | APOB | 159 | KRT1 |
| 2 | AMY2A | 81 | SERPINF1 | 160 | LMNA |
| 3 | PLG | 82 | SUPT6H | 161 | ANXA2 |
| 4 | SERPINA7 | 83 | BRD9 | 162 | DSP |
| 5 | IGF2R | 84 | APMAP | 163 | KRT9 |
| 6 | TIMP1 | 85 | TIMP2 | 164 | FLG |
| 7 | COL6A1 | 86 | SOD1 | 165 | KRT6B |
| 8 | A1BG | 87 | LGALS1 | 166 | KRT6A |
| 9 | C7 | 88 | HABP2 | 167 | FLG2 |
| 10 | HBA2; HBA1 | 89 | MRC2 | 168 | S100A7 |
| 11 | CTSD | 90 | PRDX1 | 169 | KRT10 |
| 12 | CPA6 | 91 | TPM3 | 170 | EEF2 |
| 13 | AFP | 92 | TXN | 171 | TGM3 |
| 14 | DHRS4 | 93 | TUBB | 172 | KRT17 |
| 15 | HSPG2 | 94 | PODXL | 173 | FABP5 |
| 16 | VNN2 | 95 | LGALS3BP | 174 | SFN |
| 17 | PFN1 | 96 | CLTC | 175 | FAP |
| 18 | CLSTN1 | 97 | ACTA1 | 176 | LGALS7; LGALS7B |
| 19 | QSOX1 | 98 | HSPA1B; HSPA1A | 177 | KRT80 |
| 20 | VNN1 | 99 | TRAP1 | 178 | ADAMTS13 |
| 21 | CNTN1 | 100 | LOC102723996 | 179 | CSTA |
| 22 | RGN | 101 | FLNB | 180 | ATP7B |
| 23 | H2AFZ | 102 | ALDOB | 181 | PRDX2 |
| 24 | IL1RAP | 103 | AGRN | 182 | HNRNPA1 |
| 25 | COL5A1 | 104 | FN1 | 183 | DSC1 |
| 26 | TMOD2 | 105 | PGM1 | 184 | TUBA4A |
| 27 | CCDC37; CFAP100 | 106 | HSP90AA1 | 185 | SBSN |
| 28 | SERPINA10 | 107 | GNB1 | 186 | KRT13 |
| 29 | SERPINF2 | 108 | TOP1 | 187 | KRT4 |
| 30 | AFM | 109 | HIST1H1C | 188 | ART4 |
| 31 | CHIA | 110 | LAMP2 | 189 | PZP |
| 32 | ITIH1 | 111 | HMGN2 | 190 | KRT2 |
| 33 | BMP1 | 112 | KIF2B | 191 | DSG1 |
| 34 | LTF | 113 | NCAM1 | 192 | AZGP1 |
| 35 | CD109 | 114 | ACTB | 193 | SERPINB12 |
| 36 | C5 | 115 | MFGE8 | 194 | S100A8 |
| 37 | HRNR | 116 | UBA52 | 195 | CALML5 |
| 38 | FEN1 | 117 | PGK1 | 196 | LYZ |
| 39 | POSTN | 118 | HSPA8 | 197 | S100A9 |
| 40 | SERPINC1 | 119 | PKM | 198 | DCD |
| 41 | MASP1 | 120 | GPI | 199 | HIST1H2BO |

| | | | | | |
|---|---|---|---|---|---|
| 42 | TF | 121 | GAPDH | 200 | HIST1H2BD |
| 43 | SHBG | 122 | EDIL3 | 201 | CAPN2 |
| 44 | THBS4 | 123 | MSN | 202 | HIST2H3A; HIST2H3C; HIST2H3D |
| 45 | ITIH2 | 124 | TMSB10 | 203 | H2AFX |
| 46 | CLEC3B | 125 | AHNAK | 204 | HIST1H4A; HIST1H4F; HIST1H4D; HIST1H4J; HIST2H4A; HIST2H4B; HIST1H4H; HIST1H4C; HIST4H4; HIST1H4E; HIST1H4I; HIST1H4B; HIST1H4K; HIST1H4L |
| 47 | KNG1 | 126 | ENO1 | 205 | AHSG |
| 48 | F2 | 127 | HIST1H1E | 206 | C6 |
| 49 | XPO1 | 128 | CD82 | 207 | F10 |
| 50 | LRP1 | 129 | TUBA1A | 208 | AHSG |
| 51 | ITIH3 | 130 | ARHGAP35 | 209 | DMKN |
| 52 | PLTP | 131 | VTN | 210 | APOH |
| 53 | ITIH4 | 132 | THBS1 | 211 | MT1G |
| 54 | HS6ST1 | 133 | IGFBP7 | 212 | MYADM |
| 55 | RBP4 | 134 | FSTL1 | 213 | CD47 |
| 56 | C3 | 135 | CYR61 | 214 | CD59 |
| 57 | GSN | 136 | PPIA | 215 | SLC16A3 |
| 58 | C4B; C4B_2; LOC100293534 | 137 | EEF1A1 | 216 | ATP1A1 |
| 59 | ALB | 138 | RHOC | 217 | LSR |
| 60 | HSP90B1 | 139 | CD58 | 218 | SLC1A5 |
| 61 | JUP | 140 | SLC2A1 | 219 | BASP1 |
| 62 | EFEMP1 | 141 | SLC2A14 | 220 | ITGA6 |
| 63 | MT1B | 142 | CD63 | 221 | SLC39A10 |
| 64 | CDH13 | 143 | EGFR | 222 | HLA-A |
| 65 | F13A1 | 144 | PTX3 | 223 | CDC42 |
| 66 | GC | 145 | NT5E | 224 | BSG |
| 67 | FGB | 146 | ATP2B4 | 225 | ITGA3 |
| 68 | A2M | 147 | CD44 | 226 | ITGB1 |
| 69 | LOC101929530; FANCD2P2 | 148 | LHFPL2 | 227 | ITGA2 |
| 70 | LUM | 149 | HLA-B | 228 | SLC3A2 |
| 71 | HBB | 150 | ITGB4 | 229 | CD81 |
| 72 | HGFAC | 151 | CD9 | 230 | NPTN |
| 73 | FBLN1 | 152 | C9 | 231 | RFTN1 |
| 74 | MAGEC3 | 153 | KRT78 | 232 | F3 |
| 75 | COL1A1 | 154 | CASP14 | 233 | MCAM |
| 76 | PCLO | 155 | TGM1 | 234 | ATP2B1 |
| 77 | COMP | 156 | KRT14 | 235 | SLC43A3 |
| 78 | PPARD | 157 | KRT5 | 236 | GNB2 |
| 79 | SERPIND1 | 158 | KRT16 | 237 | MARCKS |

**Table S2: Primers for qRT-PCR analysis**

| Gene | Forward Sequence (5'-3') | Reverse Sequence (5'-3') |
|---|---|---|
| *ACTA2* | GTGTTGCCCCTGAAGAGCAT | GCTGGGACATTGAAAGTCTCA |
| *CCN2* | TGGAGTTCAAGTGCCCTGAC | CTCCCACTGCTCCTAAAGCC |
| *COL1A1* | TGCTCGTGGAAATGATGGTG | CCTCGCTTTCCTTCCTCTCC |
| *GAPDH* | GCACCGTCAAGGCTGAGAAC | GCCTTCTCCATGGTGGTGAA |
| *IL6* | ACTCACCTCTTCAGAACGAATTG | CCATCTTTGGAAGGTTCAGGTTG |
| *KGF* | AGGCAGACAACAGACATGGAAT | TCGATCCTCAGGTACCACTGT |
| *MMP1* | GGGGCTTTGATGTACCCTAGC | TGTCACACGCTTTTGGGGTTT |
| *S100A10* | GGGCTTCCAGAGCTTCTTTT | CTTCTATGGGGGAAGCTGTG |
| *S100A11* | TGTCCTTGACCGCATGATGAA | TTCTGGGAAGGGACAGCCTT |
| *S100A12* | CTTCCACCAATACTCAGTTCGG | GCAATGGCTACCAGGGATATG |
| *S100A13* | TTCTTCACCTTTGCAAGGCA | GAGAGCCCACATCCTTGAGC |
| *S100A14* | CTCATGCCGAGCAACTGTG | GGGTACAGGGTGGTGGTAGA |
| *S100A16* | ATGCTGTCGGACACAGGG | TGATGCCGCCTATCAAGGTC |
| *S100A4* | TCTTGGTTTGATCCTGACTGCT | AAGCACGTGTCTGAAGGAGC |
| *S100A6* | AAGCTGCAGGATGCTGAAAT | CCCTTGAGGGCTTCATTGTA |
| *TBP* | GAGCCAAGAGTGAAGAACAGTC | GCTCCCCACCATATTCTGAATCT |
| *TUBA3C* | AGGAGTCCAGATCGGCAATG | GTCCCCACCACCAATGGTTT |
| *VEGFA* | AGGGCAGAATCATCACGAAGT | AGGGTCTCGATTGGATGGCA |
| *VIM* | AGTCCACTGAGTACCGGAGAC | CATTTCACGCATCTGGCGTTC |

**Supplementary Methods**

***Size distributions of EVs***

Samples were initially diluted at 1:20 with DPBS to achieve a concentration below $10^9$ particles/ml. Samples were introduced in the instrument using a syringe attached to a pump. For each sample, three videos were captured for 60 s with a camera level between 13 and 16. All measurements were carried out at room temperature and the chamber was cleaned with 10% ethanol and DI water between each sample. Videos were analyzed using NanoSight with detection threshold set within 5 to 10 to obtain vesicle concentration (particles/ml) and size distribution (nm).

***Tumor stiffness mapping using microindentation***

The tumor section was mounted on a customized stage and DPBS was applied to keep the tissue hydrated throughout the measurement. Dynamic indentation by nanoindenter (Nanomechanics Inc.) was used to characterize the tumor elastic modulus (Akhtar *et al.*, 2018). Sneddon's stiffness equation (Sneddon, 1965) was applied to relate dynamic stiffness of the contact to the elastic storage modulus of the samples (Herbert, Oliver and Pharr, 2008; Herbert *et al.*, 2009). 500 µm flat cylindrical probe was used in the indentation experiments. Briefly, procedure of indentation is comprised of 3 steps: 1) approaching and finding tissue surface at the indenter's resonant frequency to enhance contact sensitivity and accuracy, 2) pre-compression of 50 µm to ensure good contact, 3) dynamic measurement at 100 Hz oscillation frequency with amplitude of 250 nm. The indentation procedure mentioned above was done consecutively on multiple regions of a single tissue surface in a grid pattern to obtain stiffness map of the tumor. Because obtaining a perfectly flat tissue surface was difficult due to tissue heterogeneity, individual indentation processes were observed using a microscope camera to determine inappropriate contact of the probe to the tissue for inaccurate measurement which were excluded from data. Typically, the number of indentation points per tissue mapping was 20-40 with the resolution of 1-2 mm spacing between points depending on the size of tumor sample. The duration of stiffness mapping was 30 min on average.

***Patient tissue vesicle collection and characterization***

After collection in DPBS, primary patient tissue samples underwent bulk compression and microindentation to determine stiffness. Each sample was then transferred to 5 mL of 1% penicillin-streptomycin solution in 013-CV DMEM and incubated at 37ºC overnight. After 24 h, the supernatant of each sample was collected and the patient tissue was transferred to 120 mL of formalin and left at room temperature. The supernatants then underwent differential centrifugation to pellet out cells and cellular debris, being spun down at the following settings: 4ºC at 800 g for 5 min, 2,000 g for 10 min and 10,000 g for 30 min. The supernatants were isolated after every spin and at the end of the 10,000 g spin, they were filtered using 0.22 µm PES filters (Genesee) into an Amicon $^{R}$ Ultra - 15 10 K centrifugal filter. The Amicon spin was conducted at the setting of 4ºC at 5,000 g for 40 min, per manufacturer protocol. After the final spin, the concentrates were measured and deposited into eppendorfs. 50 µl of each sample was placed in a separate eppendorf with 950 µl of PBS, to create a sample with a dilution factor of 20. The diluted samples were then taken for nanoparticle tracking analysis and diluted further as needed.

***EV proteomics***

EVs were collected from three biological replicates of cell cultures grown on tissue culture plastic, 25 kPa, and 0.5 kPa matrices. Bligh-Dyer extraction was used to precipitate the protein content, while lipids and polar metabolites were removed in the organic (lower) and aqueous (upper phases) (Bligh, E.G. and Dyer, 1959). Briefly, 200 µL of methanol and 100 µL of chloroform were added, and the samples were vortexed for 1 min and sonicated for 5 min to break the EV structures. 100 µL of chloroform were added followed by 100 µL of water to precipitate the protein contents into the interphase. The samples were centrifuged at 14,000 x g for 10 min at room temperature. After refrigeration at -20ºC (this did not freeze either phase) overnight the precipitated proteins settled in the interphase. The methanol/water (upper) phase, and the lipid-containing chloroform (lower) phase were discarded. The remaining protein containing pellet was lyophilized to dryness and reconstituted in 20 µL of 50 mM triethyl ammonium bicarbonate pH 8.0 in 50% aqueous 2,2,2-triflouroethanol (TFE) to solubilize the precipitated proteins. The samples were reduced in 5 mM TCEP at 37ºC for 30 min and alkylated in 10 mM IAA at room temperature for 15 min in the dark. The sample was diluted 10 x in 50 mM triethyl ammonium bicarbonate (pH 7.5) and digested with 2 μg of Trypsin / Lys-C Mix at 40ºC for 16 h. Each sample was labeled with 41 µl of a different 10-plex TMT reagent for 90 min. The labeling reactions were quenched with 8 µl 5% hydroxylamine for 15 minutes, and the labeled samples were combined and lyophilized. The pooled sample was reconstituted in 2% acetonitrile, 0.1% trifluoroacetic acid and fractionated by high pH solid phase extraction on an Oasis HLB plate (Waters Corp). The peptides were eluted sequentially off the Oasis plate into separate fractions using solvents containing 5%, 10%, 25%, and 75% acetonitrile in 10 mM triethyl ammonium bicarbonate pH 8.5. The fractions were lyophilized and reconstituted in 50 μL of 0.1% formic acid 2% acetonitrile prior to duplicate LC-MS analysis.

Five microliters (10%) of each fraction were separated over a binary reversed phase gradient using aqueous 2% acetonitrile 0.1% formic acid (mobile phase A), and 0.1% formic acid in 90% acetonitrile (mobile phase B) as follows: Sample loading in 0% B with a rapid jump to 8% B, a linear ramp from 8% to 30% B over 90 min, linear ramp to 48% B over 15 min, ramp to 95% B over 5 min, 9 min hold at 95% B, and drop back to 0% B in 1 min. Data dependent acquisition DDA was used to quantify peptides and consequently proteins. The precursor scan spanning 400-1600 m/z was acquired at 120000 (at m/z = 200) resolution with automatic gain control (AGC) set to 200000 and 50 ms maximum ion injection time (IT). Precursor ions in the +2 to +6 charge states were individually isolated in 0.4 Da isolation windows and serially fragmented in order of highest intensity by high energy collisional dissociation (HCD, 38 normalized collision energy) for 4 s after each precursor scan. Fragment ions were detected from 120-2000 m/z scan at 50000 resolution, using 50000 AGC and 86 ms maximum IT. Previously fragmented ions were excluded for 30 s to prevent redundant precursor sampling.

The collected data (4 fractions analyzed in duplicate) were combined and searched against the SwissProt *Homo Sapiens* database with Mascot (v.2.6.2 Matrix Science) in Proteome Discoverer 2.4 (Thermo) using 5 ppm precursor and 0.01 Da fragment mass error tolerances, trypsin as enzyme, allowing for two missed cleavages, with cysteine carbamidomethylation and TMT on N-terminal amino acids as fixed modifications, and TMT labeling of lysine, methionine oxidation, and deamination of asparagine and glutamine as variable modifications.

Mascot ".dat" files were validated with Percolator with peptide identifications filtered at <5% FDR. The reporter ions from the Peptide Spectral Matches (PSMs) assigned to proteins were used to calculate, normalize and scale protein abundances across all samples. Relative ratios were calculated from normalized median protein abundances of biological replicates.

The heatmap was created in RStudio using R version 3.6.3 and function heatmap.2. The heatmap contains all proteins identified in the proteomics experiment. The colors correspond the abundance ratio between two EV conditions. Volcano plots, created in Thermo Proteome Discoverer 2.4.1.15, include all proteins with those proteins that are enriched 2-fold in the EV condition in the numerator in red and those enriched 2-fold in the denominator in green. To perform gene ontology, express analysis was run on those proteins enriched 2-fold or greater in Metascape.[69]
